# Wavelet for binocular vision modeling

**DOI:** 10.1101/256867

**Authors:** Louis Gaudart, Jean Gaudart

## Abstract

In the present study, binocular vision properties were modeled using a single elementary wavelet. Opponent responses (ON-OFF) appeared in the first stages of the neural coding in the retina. This property was assumed to build an adequate wavelet showing a positive part (On) and a negative part (OFF). We have examined the experimental orientation and position disparity given by Bishop. We assumed that the theoretical position disparity was given by a combination of two wavelets for a given orientation disparity β. A change in β implied a change in the magnitude of one of the wavelets and consequently a change in the wavelets combination. There was a close match between the theoretical and experimental position disparity curves according to the changes in orientation disparity.

## 1. Introduction

Several previous physiological studies have shown that there are various receptive field organizations within the retina, the lateral geniculate nucleus and the cortical level [37]. The type 1 exhibits center-surround spatial and chromatic opponency. The type 2 exhibits a chromatic opponency (*e.g*. blue-yellow), without spatial opponency. The modified type 2 exhibits a center-surround spatial opponency and a chromatic opponency only in the center. The type 3 exhibits an achromatic center opposed by an achromatic surround. The double opponent cells (DOC) [1] exhibit a center-surround organization with a chromatic opponency in the center (for example R+G-) and an opposed chromatic opponency in the surround (for example R-G+).

Many attempts have been made to describe the physiological properties of the visual system with the help of various theoretical models. Rodieck [33] and then Enroth-Cugell and Robson [12] introduced a difference of two Gaussians (DOG) to model ganglion cell receptive fields. Another model was given by Marcelja [27] and Daugman [8] who introduced the Gabor function and the joint space/spatial frequency uncertainty relation. The Gabor transform provides a local description of the visual process. However, the Gabor model is inadequate in some species [35, 39, 18]. Marr [28] used the Laplacian of the Gaussian (Δ^2^G) called the “Mexican hat”. However, this author had outlined that this function cannot describe a type 1 chromatic receptive field [29]. Kulikowski, Marcelja, and Bishop [20] used a Gabor function to relate the receptive fields to the spatial frequency and orientation. Billock [1] reported a filtering mechanism to model the double opponent receptive field. Croner and Kaplan [6] investigated the achromatic receptive field (type 3) and contrast responses of ganglion cells using the difference of two Gaussians.

In a more general way, the previous theories were made under various hypotheses but failed to explain several visual properties simultaneously. Most problems were considered individually, in isolation from other visual process. The above theoretical studies examined directly the spatial receptive field organization. However, another possible type of model can be obtained by investigating the first stages of the visual process, from visual receptors to ganglion cells. Chromatic and achromatic signals are carried by several elementary cells as cones, horizontal, bipolar, amacrine, or ganglion cells [21]. A receptive field structure is the result of several specific connectivities of these neurons. In order to explain color vision and light adaptation, Guth [15, 16], then De Valois and De Valois [9, 10] proposed multi-stage models.

Using a multiresolution approach for a vision process model is attractive and remains an open question. Wavelets are mathematical functions which are localized in space and analyze the signal according to scale. A sharp or a coarse analysis is known as a multiresolution analysis. The wavelet theory can be used in two different manners. First, the wavelets can be considered as a mathematical tool to investigate a signal as an electroencephalogram one [32, 34] or an electrocardiogram one [36]. The wavelet theory is especially used to investigate digital image coding [24, 25, 17, 30]. Second, the wavelets can be considered as a mathematical tool for modeling physiological properties. Li and Atick [23] have shown the principle of efficient visual coding as implemented through multiscale representation. Bradley [3] described a model of the human visual system based on the wavelet transform.

The choice of an adequate wavelet function is always a crucial and difficult factor. To build an adequate wavelet we have to investigate the first stages of the neural coding. Numerous studies [22, 38, 7] have shown that the opponent responses (ON-OFF responses) appear in the early neural synapses. An adequate wavelet must take into account this property. Moreover, considering the binocular vision, the visual judgments of the depth produce by the disparity differences of two vertical lines play an important role [40, 26]. The studies of Bishop [2] concerning the position and orientation disparities are particularly interesting. This author showed how signals originating in each eye are combined to obtain a binocular standard receptive field. Therefore, these results have to be taken into account to build a mathematical model.

We have to address the problem from the following standpoint. Is it possible to have a single conceptual framework to provide a description of several processes? In such a way, we built a wavelet which verified the three following conditions:

a. The theoretical curves obtained have to be similar to corresponding experimental curves ON-OFF response and Bishop data [2]).
b. The number of required theoretical coefficients have to be small and each of these have correspond to a given property of the visual system.
c. The model can be used to account other visual mechanisms.

## 2 Methods

### Wavelet function

The basic characteristics of the early visual processing, inside the retina, are ON-OFF responses. There are parallel pathways through the retina, with On-channels and Off-channels [19, 40]. A receptive field is composed of an ON-region and an OFF-region. The simplest function to model an “ON-OFF” signal is the Haar function H, defined by the following relations [4]:

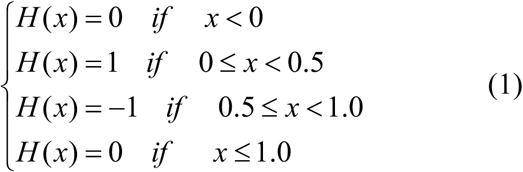

However, the Haar function, which is a wavelet, is not an adequate model to reflect a visual processing. Indeed, this function shows angular points with a gap between the positive and the negative value, whereas the response of the visual system is continuous and regular, such as the transmission of a signal by a receptive field. A more appropriate wavelet should be determined. A continuous rising or decaying phase of a curve can be modeled by hyperbolic functions. We have defined over ***R***\* a function Ψ, having a positive and negative region, by:

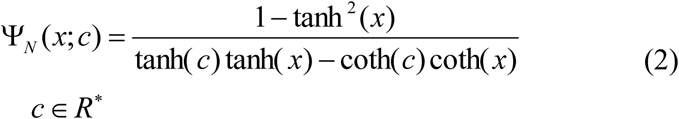

Such a curve can have a shape similar to a Haar curve (Fig. 1).

**Fig. 1.**
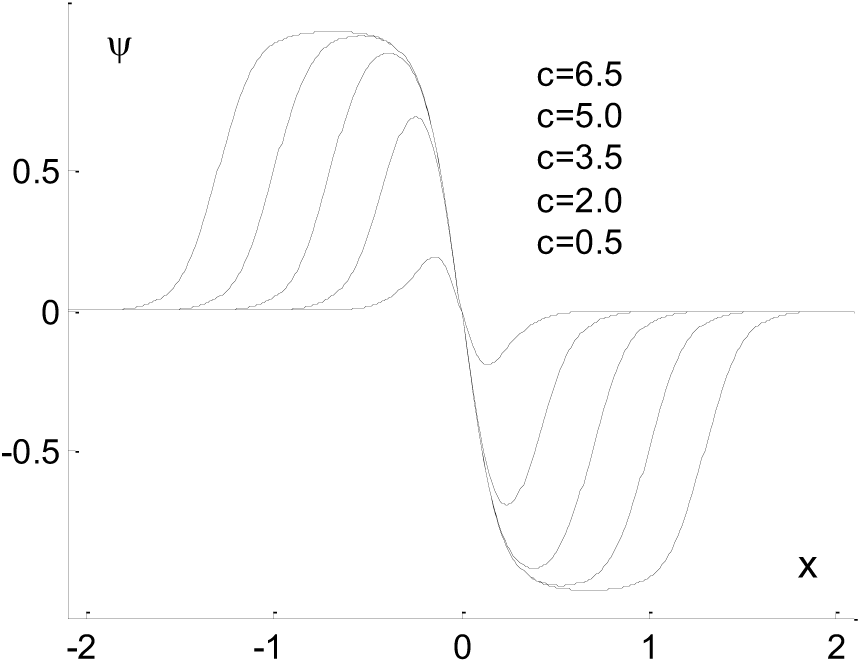
Plots of the function Ψ, for five c-values. For all curves, the coefficients “a” and “z” are the same: a = 0.2; z = 0.

We have checked that the function Ψ Wverifies the following conditions [4].

a. The function integral converges: 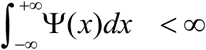
b. The function integral is null: 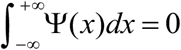
c. The Fourier transform of the function is null when *ω* = *0*.
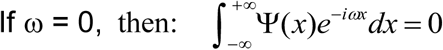
d. The function has a higher number of vanishing moments (1-order moment, 2-order moment, the even moments are null).

Consequently, the function Ψ is a wavelet, assuming a positive and a negative part. Thus, it is particularly well adapted to reflect an ON-OFF signal.

### Wavelet family

The function specified by the above formula (Eq. 2) is a “mother wavelet”. From the mother wavelet we have obtained a wavelet family by introducing a dilation coefficient “a” and a translation coefficient “z”. Therefore, the family wavelet is given by the following formula:

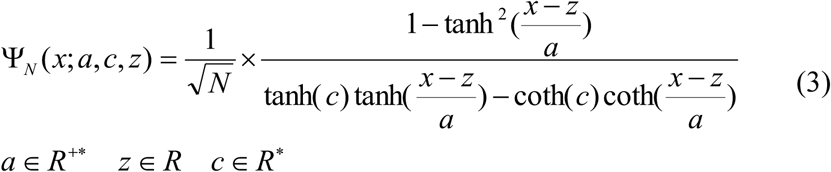

Note that the function contains the coefficient “c” which does not form part of the classical wavelet theory. However, this coefficient plays an important role, as we will see further on. It allows taking into account some of physiological properties of the visual system, inducing the shape of the curve (Fig. 1). If “c” increases, then the curve approaches a Haar function. For a given value of “c”, the theoretical coefficient “a” determines the wavelet width (Fig. 2). By varying the translation coefficient “z”, one can scan the x-space (Fig. 3).

**Fig. 2.**
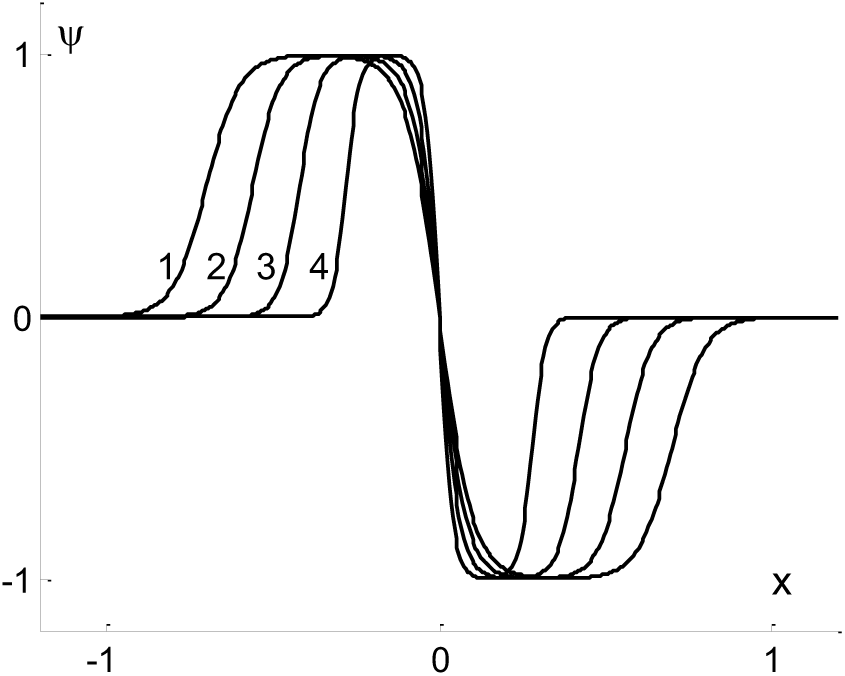
Plots of the function Ψ, for four a-values. Curve 1: a = 0.10; curve 2: a = 0.08; curve 3: a = 0.06; curve 4: a = 0.04. For all curves, the coefficients c and z are the same: c = 7; z = 0.

**Fig. 3.**
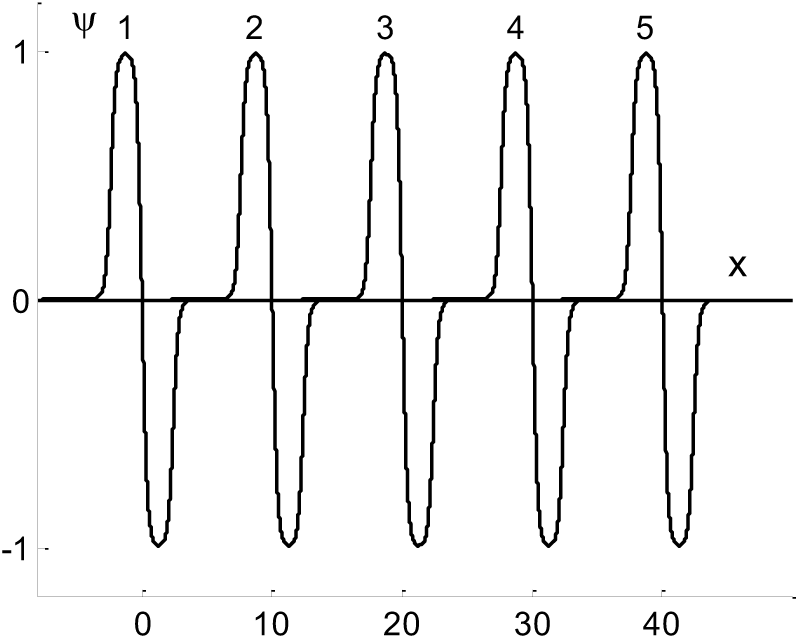
Plots of the function Ψ, for five z-values. Curve 1: z = 0; curve 2: z = 10; curve 3: z = 20; curve 4: z = 30; curve 5: z = 40. For all curves, the coefficients a and c are the same: a = 0.4; c = 6.

### Bishop’s data

We compared the above wavelet versus the data published by Bishop [2]. The neural mechanisms have been studied by this author in the striate cortex of the cat. A stereopair of lines was viewed in a stereoscope. The two retinal images, XRYR (right eye) and XLYL (left eye) were at an angle to one another. The position disparity was deduced to the length of the line YLYR. The angle between the two images was the orientation disparity. The orientation and position disparity changes occurred in the right eye with the left eye image held constant. The position and orientation disparities were expressed in terms of a visual angle. The data given by Bishop [2] are represented by the points in Fig. 5a and 5b. The angle β was the orientation disparity for the right eye. The left eye image held constant at 93°. The data points were recorded in orientation steps of 3°.

**Fig. 5a and 5b.**
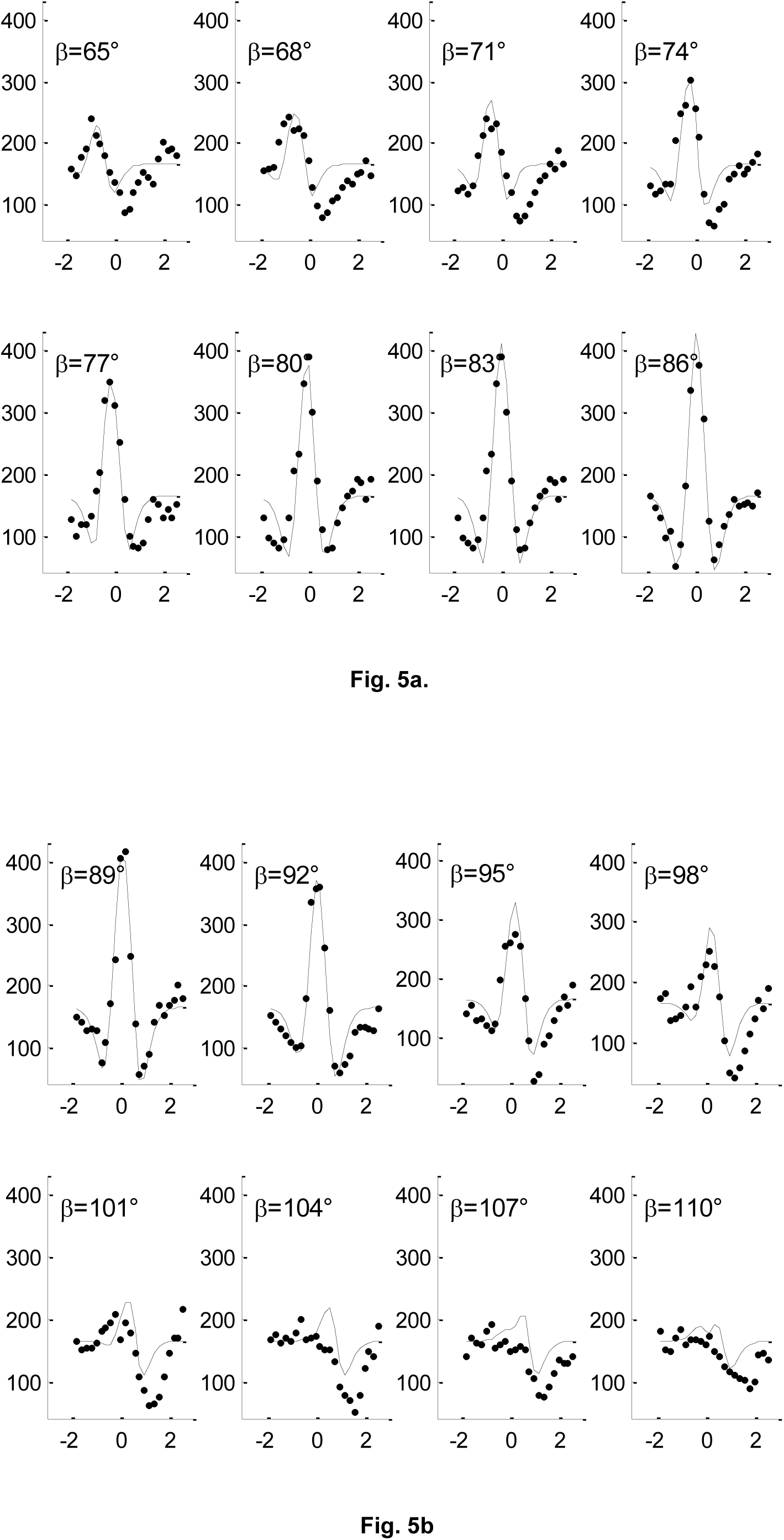
At each orientation disparity β, a curve of position disparity is plotted (x-axis in degrees). The points represent the position disparity redrawn from Bishop [2]. The y-axis represents the total spike count. The orientation disparities β are given at increment of 3°. The continuous lines are obtained by combining two wavelets according to Eq. 5.

## 3 Results

If we assumed that one retina response is modeled by a wavelet, then, the combination of two wavelets was able to reproduce the position and orientation disparities (two retina responses).

First, let us consider two wavelets (Fig. 4) defined by:

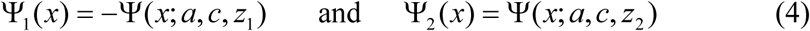

**Fig. 4.**
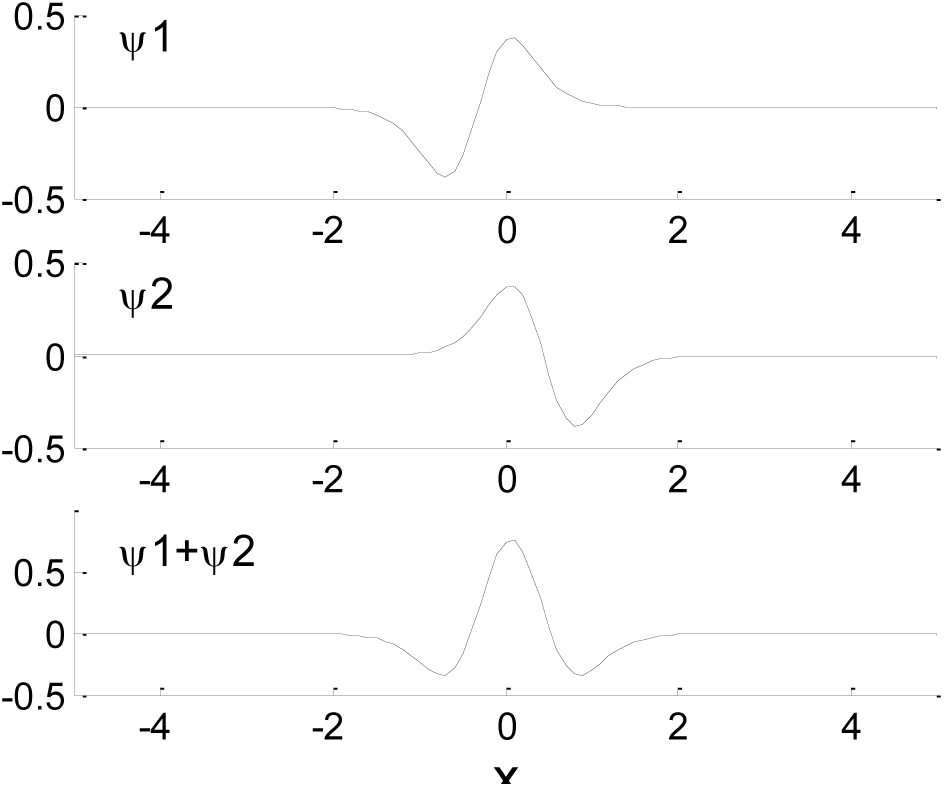
The upper figure shows the wavelet Ψ1(x;a, c, z1), the central figure shows the waveletΨ2(x;a, c, z2) and the lower figure shows a binocular response. The wavelet coefficients are: a = 0.46; c = 1; z1 = - 0.32; z2 = 0.44. The three curves share a common abscissa.

We introduced a theoretical angle α to represent the orientation disparity β and two parameters A and B. The full amplitude of the resulting theoretical curve was adjusted to the corresponding Bishop’s curve by varying the parameter A and the position of the resulting theoretical curve was given by the parameter B. Thus, we obtained the following equation:

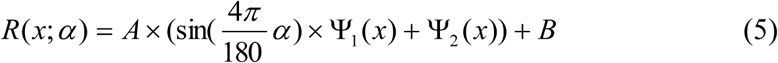

where x is the position disparity expressed in degree.R(x;α) represents the theoretical binocular response for a given theoretical orientation disparity α.

The locations of wavelet centers are z1 and z2, respectively. The distance between the two centers represents the distance between the two retinas. During the experimental measurements this distance was held constant: z2 - z1 = constant. Second, the calibration of the model was achieved through a middle value of the orientation disparity: β = 86°. For this value the experimental curve was a standard receptive field. To model this curve, we have determined the values of the wavelet coefficients. The coefficient “c” was chosen in an arbitrary manner by setting: c = 1. The other coefficients were calculated by minimizing the normalized root mean squared deviations (NRMSD) between the data and the theoretical values. We obtained: a = 0.46, z1 = - 0.32, A = 360, B = 165.

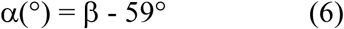

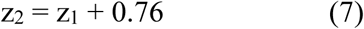

The parameter A converts an arbitrary theoretical unit into the unit of Bishop’s data (a number of spikes).

By using Eq. 5 we obtained the continuous curve of the Fig. 5a for β = 86°.

Third, the following results were obtained by using Eqs 5, 6 and 7, and the above values of “a”, A, and B. The disparity theoretical curves for the others β-values were performed by varying α. As mentioned above, an increase in coefficient “c” produced an increase in the theoretical curves. To match the theoretical curve to the experimental points, we calculated a value of the coefficient “c” for each β-value. Following the same idea we calculated a z1-value for each β-value.

Table 1 summarizes the performance of the coefficients calculations. The NRMSD given in this table are the minimal values which were obtained by varying “c” and “z1”.

**Table 1.** Wavelet coefficients “c” and “z1” for a given orientation disparity.

| $\beta$ | c | z<sub>1</sub> | NRMSD* |
| --- | --- | --- | --- |
| 65° | 0.35 | - 1.14 | 0.26 |
| 68° | 0.40 | - 0.94 | 0.28 |
| 71° | 0.45 | - 0.84 | 0.23 |
| 74° | 0.55 | - 0.64 | 0.17 |
| 77° | 0.70 | - 0.54 | 0.13 |
| 80° | 0.80 | - 0.45 | 0.11 |
| 83° | 0.90 | - 0.38 | 0.06 |
| 86° | 1.00 | - 0.32 | 0.04 |
| 89° | 1.00 | - 0.29 | 0.06 |
| 92° | 0.90 | - 0.30 | 0.08 |
| 95° | 0.76 | - 0.21 | 0.13 |
| 98° | 0.66 | - 0.14 | 0.16 |
| 101° | 0.40 | - 0.12 | 0.33 |
| 104° | 0.40 | + 0.11 | 0.36 |
| 107° | 0.39 | - 0.05 | 0.34 |
| 110° | 0.32 | - 0.01 | 0.43 |

\* Normalized Root Mean Squared Deviation.

* Normalized Root Mean Squared Deviation.

On the basis of this analysis, the theoretical curves of the Fig. 5a and Fig. 5b were obtained. The relationship between “c” and the orientation disparity is revealed by the Fig. 6. To a first approximation, two linear relations can be regarded as satisfactory.

## 4 Discussion

We defined a wavelet function which take into account different processes of the visual information. We can obtain a plausible model by considering a certain number of visual properties.

### Shape of the theoretical and experimental curves

We now have good evidence that there are ON-OFF responses in the earlier levels of the retina [31]. Consequently, we defined a wavelet which presents a positive region (ON), and a negative region (OFF). Then, we could expect that such wavelet was able to model binocular vision properties. Indeed, a good agreement was shown with the data found by Bishop [2], in particular with the position and orientation disparities. It is particularly interesting to compare the theoretical curves and the experimental points of the Fig. 5a and Fig. 5b. The theoretical changes of the curves were obtained by varying the theoretical orientation disparity α (Eq. 5). Likewise, the a-change was deduced from the p-change using Eq. 6. On the other hand, it is interesting to note that both theoretical and experimental curve changes are the same. We can see figure 5a, for β = 65°, is a unique right-hand minimum for a positive position disparity. As β (or α) increases, a second left-hand minimum appears, located in a negative position disparity. This minimum increases until the value of β reaches 86°. As β still increases, the second minimum progressively decreases and disappears about β = 104°.

### Theoretical coefficients

The required coefficients were in few in number: “c”, “z”, and “a”. These estimated coefficients have significant physiological interpretations.

Let us recall that the coefficient “c” was not deduced from the wavelet theory. Nevertheless it has a specific meaning in the case of the binocular vision. Bishop [2] noticed that any binocular response was larger than the sum of the corresponding monocular responses. This property is reflected by the coefficient “c”. The visual information processing achieved through different channels. A channel feature should be characterized by a value of “c” which is similar to an amplification coefficient. The Fig. 6 shows that the coefficient “c” is not an arbitrary coefficient. Indeed, the coefficient “c” can be deduced from the β-value.

**Fig. 6.**
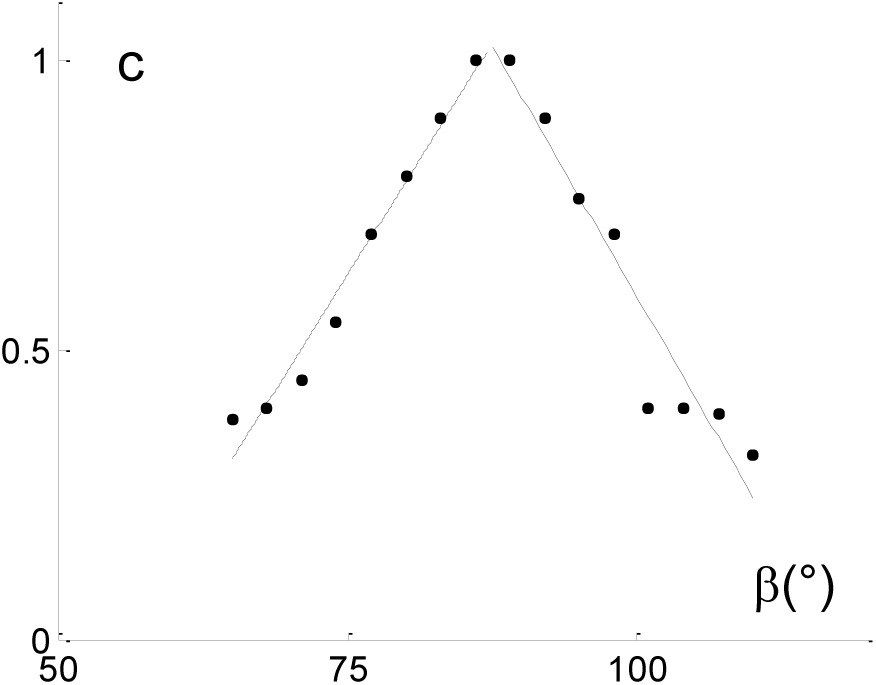
Coefficient “c” change plotted as a function of orientation disparity β. The points were deduced from the Table 1. The straight lines were obtained by use of a linear regression.

The coefficient “a” determines the wavelet width. The visual system adapts to the image size. It can see a large surface or a small point. This property is reflected by the dilation coefficient. The smaller the wavelet step, the better the analysis. The Bishop’s data were obtained with a stimulus width held constant. Consequently, the dilation coefficient did not play any role in the present study. The representation of an image in the visual cortex involves both spatial position and spatial frequency variables [20]. In the present study the spatial variable is the position disparity x and the spatial frequency variable is the coefficient “a” as we deduced from the Eq. 3.

The coefficient “z” is the classical translation coefficient of the wavelet theory. In the present study it indicates the spatial position of a retina with regard to the incident stimulus. If an eye position “z1” is used as a reference value, then the second eye position “z2” may be known. Moreover, the visual system is able to scan the viewed surface. This property is reflected by the translation coefficient. The actual eye movements explore a viewed surface using numerous successive saccades [5, 11]. Each saccade is reflected by a small variation Δz of the translation coefficient.

We can conclude that the estimated coefficients used in our study are consistent with the basic properties of the visual system.

### Chromatic receptive fields and contrast sensitivity function

The present wavelet model can be used to build various chromatic fields, as we have shown in a previous study [14]. We have defined chromatic wavelets in the following way. For example, the positive segment of a chromatic wavelet represented a response of a green stimulus and the negative segment represented a response of a red stimulus. Thus, we obtained a chromatic wavelet Green-Red. Such a wavelet modeled one of the curves given by De Valois and De Valois [9, 10]. Under the same conditions we obtained a Blue-Yellow wavelet. The De Valois and De Valois [9, 10] chromatic curves were obtained by combining several chromatic wavelets. Then, the simple chromatic receptive fields (type 1 and 3, for example) were obtained by a linear combination of two chromatic wavelets. The more complex chromatic receptive fields (type 2, 2-modified, Double Opponent Cells) were obtained by combining several chromatic wavelets.

After obtaining an adequate wavelet function, it becomes possible to use all the properties of the wavelet theory. The wavelet transform is a particularly appropriate tool to model other visual system properties, such as the contrast sensitivity function, or the view of an illuminance gradient between a bright and a dark area [13]. The wavelet transform provides a flexible window which automatically narrows when observing high frequency phenomena and widens when observing low frequency phenomena.

## 5 Conclusion

In the present approach, several properties of the visual system were modeled within a single conceptual framework. Even in view of the progress of wavelet analysis in recent years, the question is far from being settled. The place of the wavelet in the visual processing levels cannot easily be found. However, the wavelet function and the wavelet transform reflect a visual processing hierarchy. Such a representation is very economical in terms of coefficient number. We have shown that a model based on the wavelet theory enables one to find several visual properties. At the output of the retina, *i.e*. the ganglion cells, the wavelet is consistent with the measured responses. At another stage of the process, linear combinations are consistent with binocular vision properties. Finally, a wavelet transform can be situated at the cortical regions. We have explored the theory in simple visual systems with a static object, without eye movement. The model presented here is a first approach. Predictions of visual performances would require a study of various variables as time or movement. A more elaborated model should include recognition of objects at a distance.

